## Supplemental data for "MagIC-Cryo-EM: Structural determination on magnetic beads for scarce macromolecules in heterogeneous samples"

**A**

| | Test 1 | Test 2 | Test 3 | Test 4 | Test 5 | Test 6 | Test 7 | Test 8 | Test 9 (Shown in Fig. 2) |
|---|---|---|---|---|---|---|---|---|---|
| Input sample | H1.8-GFP nucleosome | H1.8-GFP nucleosome | H1.8-GFP nucleosome | H1.8-GFP nucleosome | H1.8-GFP nucleosome | H1.8-GFP nucleosome | H1.8-GFP nucleosome + nucleosome | H1.8-GFP nucleosome + nucleosome | H1.8-GFP nucleosome + nucleosome |
| Inner spacer module | 11 nm 3HB | 60 nm SAH | 60 nm SAH | 11 nm 3HB | 11 nm 3HB | 11 nm 3HB | 11 nm 3HB | 11 nm 3HB | 11 nm 3HB |
| Outer spacer module | 30 nm SAH | Not used | Not used | 60 nm SAH | 60 nm SAH | 90 nm SAH | 60 nm SAH | 60 nm SAH | 60 nm SAH |
| Target capturing module | Tandem GFP nanobody | GFP nanobody (LaG94-10) | GFP nanobody (LaG94-10) | GFP nanobody (LaG94-10) | GFP nanobody (LaG94-10) | GFP nanobody (LaG94-10) | GFP nanobody (GFP enhancer) | GFP nanobody (GFP enhancer) | GFP nanobody (GFP enhancer) |
| Grid | Quantifoil gold R1.2/1.3 + graphene | Quantifoil gold R1.2/1.3 + graphene | Quantifoil gold R1.2/1.3 + graphene | Quantifoil gold R1.2/1.3 + graphene | Quantifoil gold R1.2/1.3 + graphene | Quantifoil gold R1.2/1.3 + graphene | Quantifoil copper R1.2/1.3 + graphene | Quantifoil gold R1.2/1.3 + graphene | Quantifoil gold R1.2/1.3 + graphene |
| Microscope | Talos Arctica 200 kV | Talos Arctica 200 kV | Titan Krios 300 kV | Talos Arctica 200 kV | Titan Krios 300 kV | Titan Krios 300 kV | Talos Arctica 200 kV | Talos Arctica 200 kV | Titan Krios 300 kV |
| Detector | K2 (Gatan) | K2 (Gatan) | K2 (Gatan) | K2 (Gatan) | K3 (Gatan) | K3 (Gatan) | K2 (Gatan) | K2 (Gatan) | K3 (Gatan) |
| Beads | 11 nm 3HB 30 nm SAH spacer | 60 nm SAH spacer | 60 nm SAH spacer | 11 nm 3HB 60 nm SAH | 11 nm 3HB 60 nm SAH | 11 nm 3HB 90 nm SAH | 11 nm 3HB 60 nm SAH | 11 nm 3HB 60 nm SAH | 11 nm 3HB 60 nm SAH |
| 3D structure | 7.3 Å | 4.5 Å | 9.8 Å | 4.6 Å | 4.1 Å | 4.0 Å | 8.4 Å | 4.7 Å | 3.8 Å 3.6 Å (Polished) |

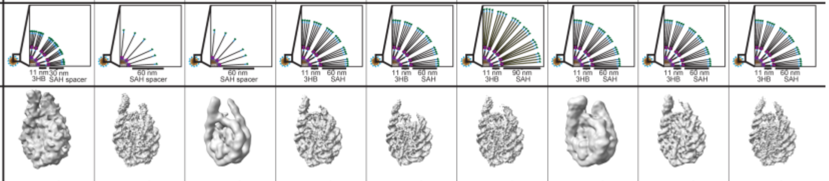

**B**

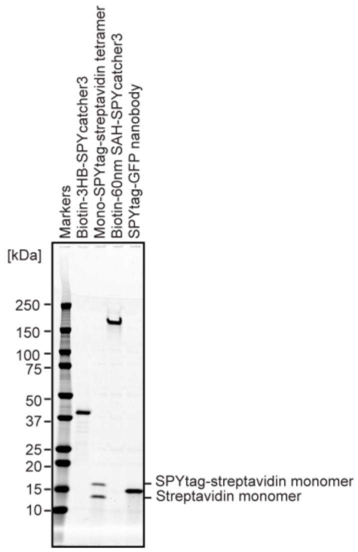

**Figure S1. Optimization of the MagIC-cryo-EM beads.**

**(A)** Spacer modules attached to the 50-nm magnetic beads were optimized to cover intense halo-like noises formed by the beads. The *in vitro* reconstituted H1.8-GFP bound nucleosomes were used for MagIC-cryo-EM optimization with various versions of the spacer modules. The cartoons depict the beads and spacer length of each experiment. The critical parameters are colored with red. The bottom 3D maps are the cryo-EM structures determined in each experiment. For the sub-5 Å resolution structure determinations using the 300 kV microscope, layers of the of 11 nm 3HB spacer and 60 nm SAH spacer are required on the paramagnetic nanobeads (Test 1, 3 and 5). For the sub-5 Å resolution structure determinations using the 200 kV microscope, the inner layers with the 11 nm 3HB spacer and mono-SPYtag avidin tetramer can be omitted because the noise signals are weaker in the 200 kV microscope then that in 300 kV microscope (Test 2 and 3). **(B)** Purified proteins for assembling the MagIC-cryo-EM beads. SDS-PAGE analysis was done by applying samples to 4–20 % Criterion TGX Precast Midi Protein Gel (BioRad 5671095) and ran at 200 V for 40 min. GelCode Blue stained gel is shown.

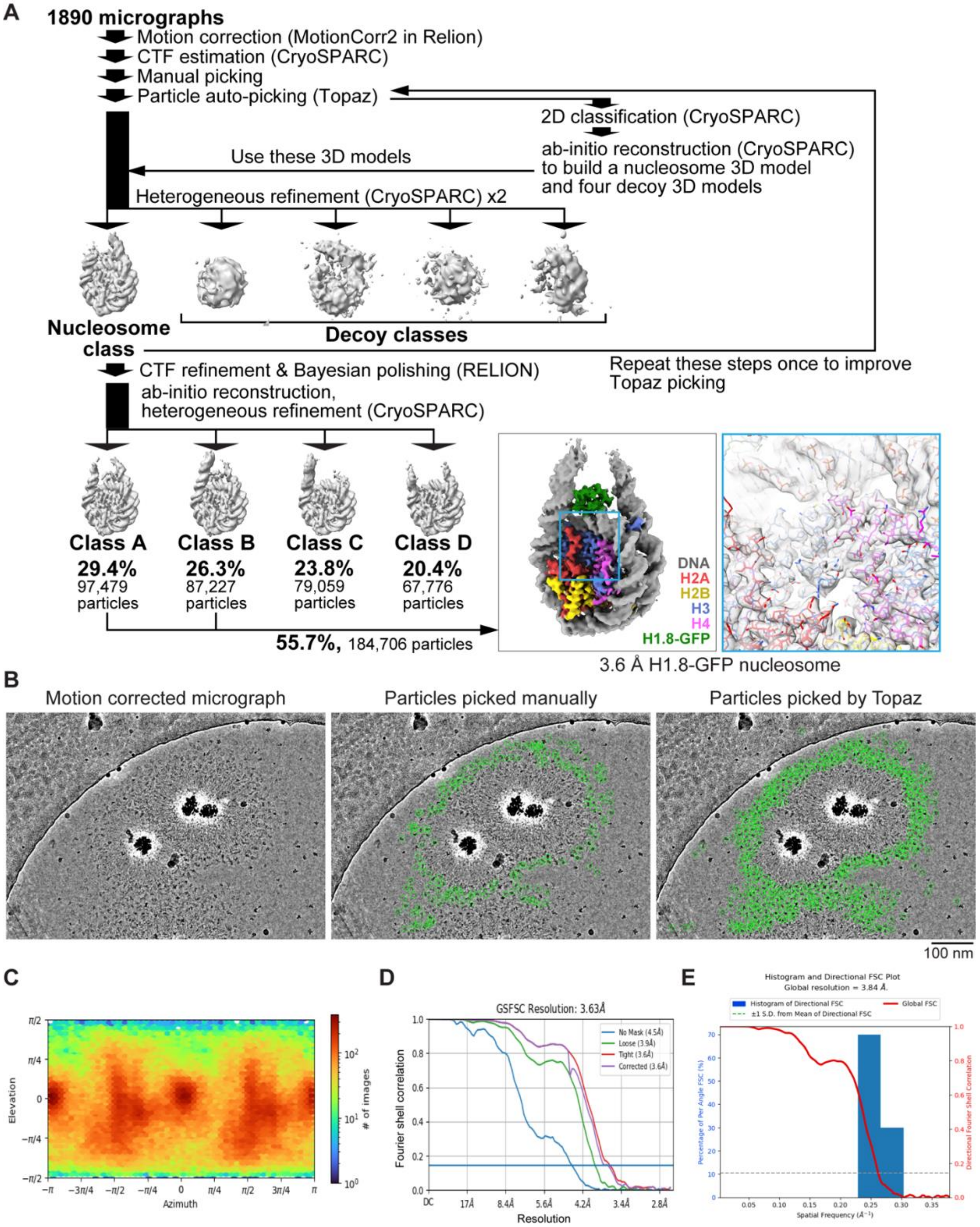

**Figure S2. MagIC-cryo-EM single particle analysis of *in vitro* reconstituted H1.8-GFP bound nucleosome. (A)** The single particle analysis pipeline for the MagIC-cryo-EM of the *in vitro* reconstituted H1.8-GFP-bound nucleosomes. Non-nucleosome or noisy particles were removed by heterogeneous refinement with decoy 3D classes (decoy classification). Using the particles assigned to the nucleosome class, another round of heterogeneous refinement was performed to isolate the classes with apparent

H1.8 densities. The particles assigned to the classes A and B were mixed, and the 3D structure of the H1.8-GFP-bound nucleosome was determined at 3.63 Å resolution. **(B)** Comparison between manually picked particles used to train Topaz (middle panel) and the particles picked by Topaz (right panel). Green circles indicate the picked particles. **(C)** Particle orientation of cryo-EM structure of the H1.8-GFP bound nucleosome. **(D)** Gold-standard Fourier Shell Correlation (FSC) curve of the H1.8-GFP nucleosome. The final resolutions of the cryo-EM maps were determined by the gold-standard with a threshold of 0.143. **(E)** 3D FSC curve of the H1.8-GFP nucleosome.

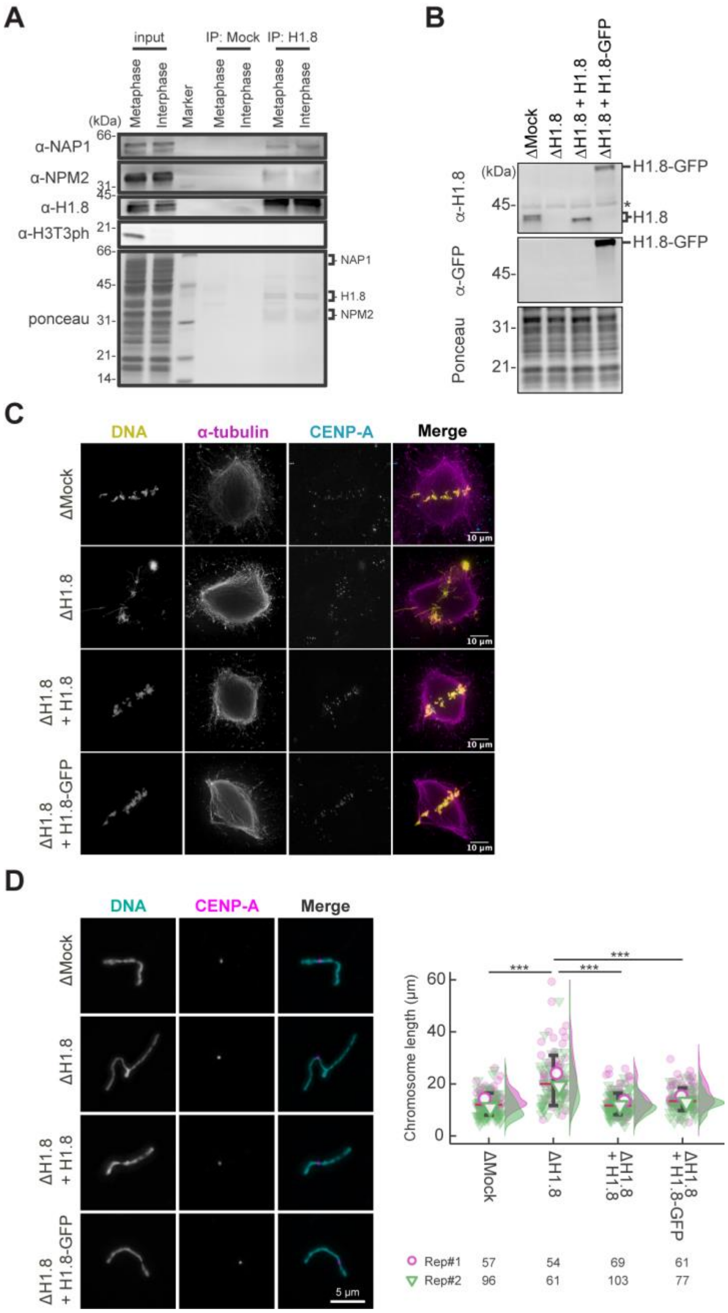

**Figure S3. Functional assessment of H1.8-GFP in *Xenopus* egg extract. (A)** Cell cycle-independent H1.8 binding to NAP1 and NPM2 in the *Xenopus* egg cytoplasm. H1.8 was immunoprecipitated from *Xenopus* CSF (metaphase) extracts or interphase extracts and analyzed by western blotting. Antibodies against phosphorylated histone H3 Thr3 H3T3ph) were used as a marker for M phase. Amounts of NAP1 and NPM2 co-immunoprecipitated with anti-H1.8 antibodies did not change between metaphase and interphase. An example of two reproducible results is shown. **(B)** Western blots to show the depletion efficiency of the endogenous H1.8 and complementation of recombinant non-tag H1.8 and H1.8-GFP in *Xenopus* egg extract. The asterisk indicates a non-specific cross-reacting band. **(C)** Representative fluorescence images of metaphase chromosomes with spindles in Mock- (ΔMock), endogenous H1.8-depleted *Xenopus* egg extract (ΔH1.8), and recombinant H1.8 or recombinant H1.8-GFP supplemented endogenous H1.8-depleted *Xenopus* egg extracts (ΔH1.8+H1.8 or ΔH1.8+H1.8-GFP, respectively). Misaligned metaphase chromosome phenotype caused by H1.8 depletion was rescued in H1.8 or H1.8-GFP supplemented *Xenopus* egg extracts. **(D)** Left; representative fluorescence images of individualized metaphase chromosomes. Elongated chromosome morphology caused by H1.8 depletion was rescued in H1.8 or H1.8-GFP supplemented *Xenopus* egg extracts. Scale bar, 5 µm. Right; quantification of chromosome length visualized by SuperPlots. Data distribution of the length of each individual chromosome from two biological replicates (purple and green) is shown as jitter plot with half violin plot. Each mark (purple open circle and green open inverted triangle) represents the average length of chromosomes from a single replicate. Bar represents median (red) and SD (black) of two biological replicates. Given the result that dataset is not normal distribution, confirmed by Shapiro-Wilk normality test, each p-value was calculated using Welch's t-test. ***, $p < 0.001$. The number of individualized chromosomes analyzed in each condition for each replicate is indicated at the bottom of the figure.

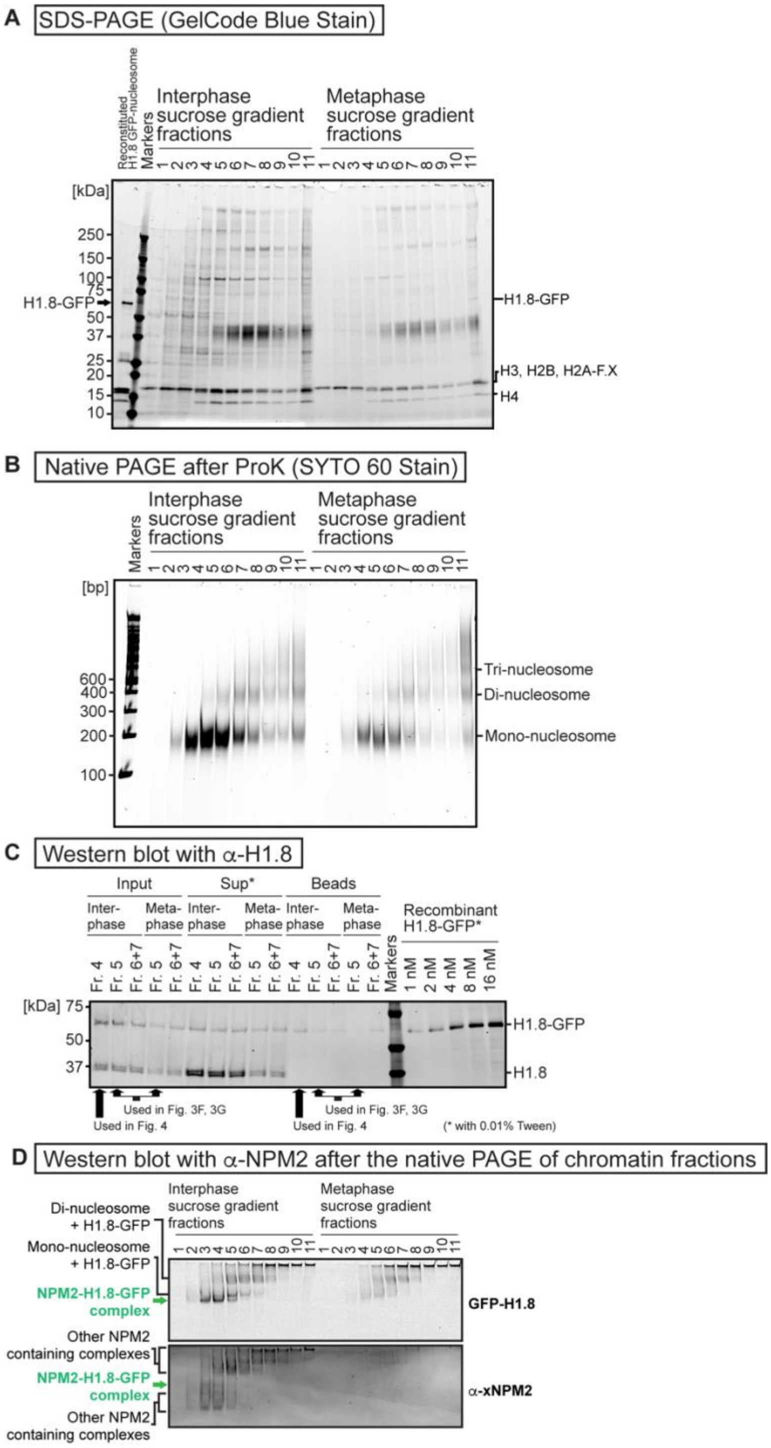

**Figure S4. Sucrose gradient fractions of the fragmented interphase and metaphase chromosomes formed in *Xenopus* egg extract. (A)** Sucrose density gradient centrifugation fractionations of MNase-treated interphase and metaphase chromosomes formed in *Xenopus* egg extract. SDS-PAGE analysis was done by applying 15 µL samples to 4–20 % Criterion TGX Precast Midi Protein Gel (BioRad 5671095) and ran at 200 V for 40 min. GelCode Blue stained gel is shown. **(B)** Native PAGE analysis of DNA lengths of the sucrose gradient fractions shown in A. 15 µL samples were treated with protease K and RNaseA and applied to 6 % native PAGE gel with x0.5 TBE and ran at 150 V for 80 minutes. **(C)** Western blot of the sucrose gradient fractions used for the MagIC-cryo-EM. H1.8-GFP and endogenous H1.8 were detected by antibody for H1.8. 'Input' lanes indicate the sucrose gradient fractions before isolating H1.8-GFP-containing complexes with MagIC-cryo-beads. 'Sup' lanes indicate the unbound fractions after the isolation by MagIC-cryo-beads. 'Beads' lanes indicate the MagIC-cryo-beads isolating H1.8-GFP-containing complexes. 'Recombinant H1.8-GFP' lanes indicate purified H1.8-GFP with known concentrations. The samples for 'Sup' and 'Recombinant H1.8-GFP' lanes contained 0.01 % Tween20, which increased the western blot signals presumably by reducing the sample absorption to the tubes and tips side wall. **(D)** Comparison of the H1-GFP and NPM2 containing bands in native PAGE of the sucrose density gradient centrifugation fractionation of MNase-treated interphase and metaphase chromosomes formed in *Xenopus* egg extract. H1-GFP was detected by GFP fluorescence (identical image with Fig 3C). NPM2 was detected by an anti-NPM2 antibody through western blotting, following protein transfer from the native PAGE gel to the membrane.

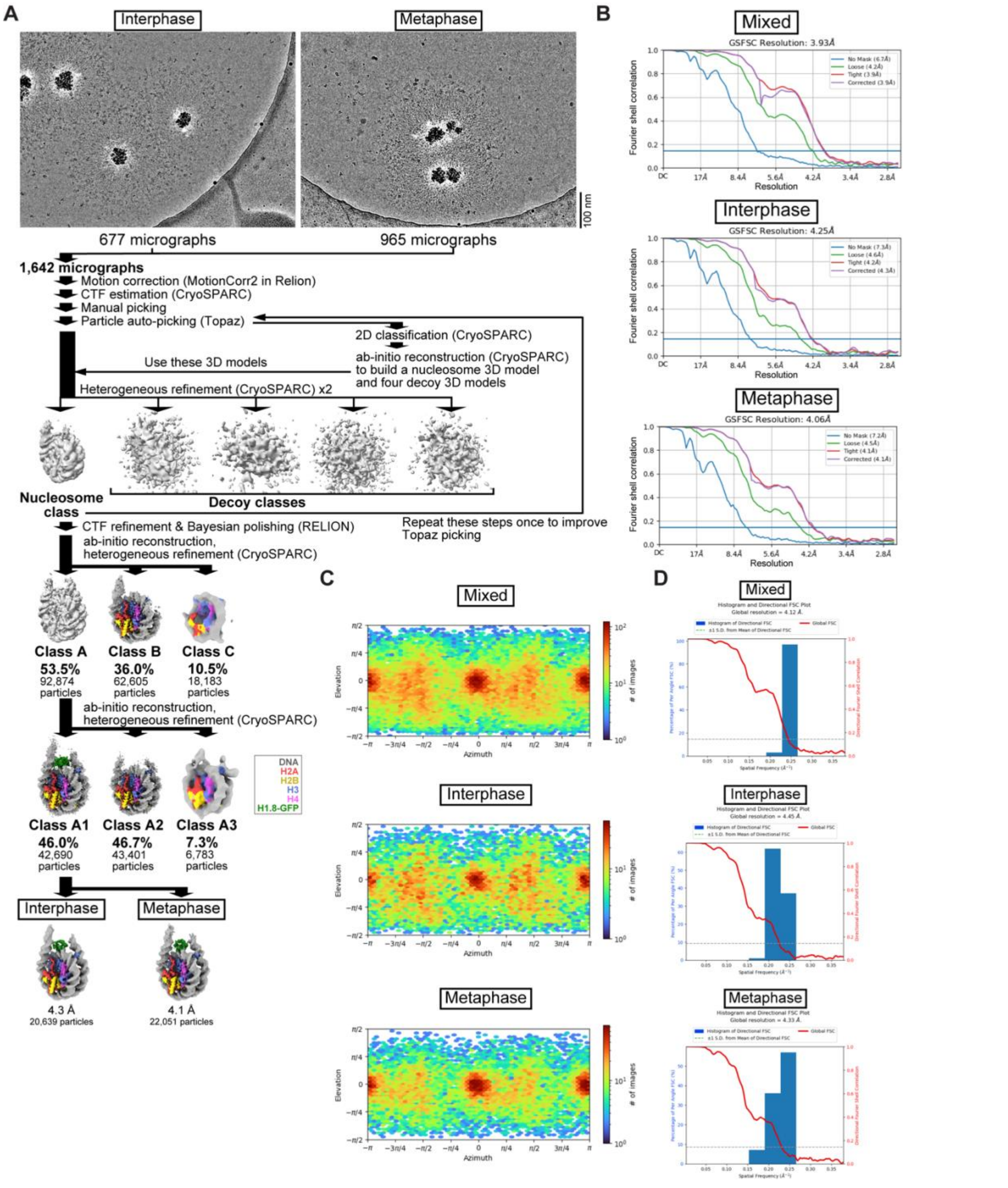

**Figure S5. Single particle analysis pipeline for the MagIC-cryo-EM of the interphase and metaphase H1.8-GFP bound nucleosomes formed in *Xenopus* egg extract. Related to Figure 3. (A)** Single particle analysis and in silico mixing 3D classification pipeline for the MagIC-cryo-EM. **(B)** Gold-standard FSC curves of the interphase and metaphase H1.8-GFP-bound nucleosomes. The final resolutions of the cryo-EM maps were determined by the gold-standard with a threshold of 0.143 **(C)** Particle orientation of cryo-EM structure of the interphase and metaphase H1.8-GFP-bound nucleosomes. **(D)** 3D FSC of the interphase and metaphase H1.8-GFP-bound nucleosomes.

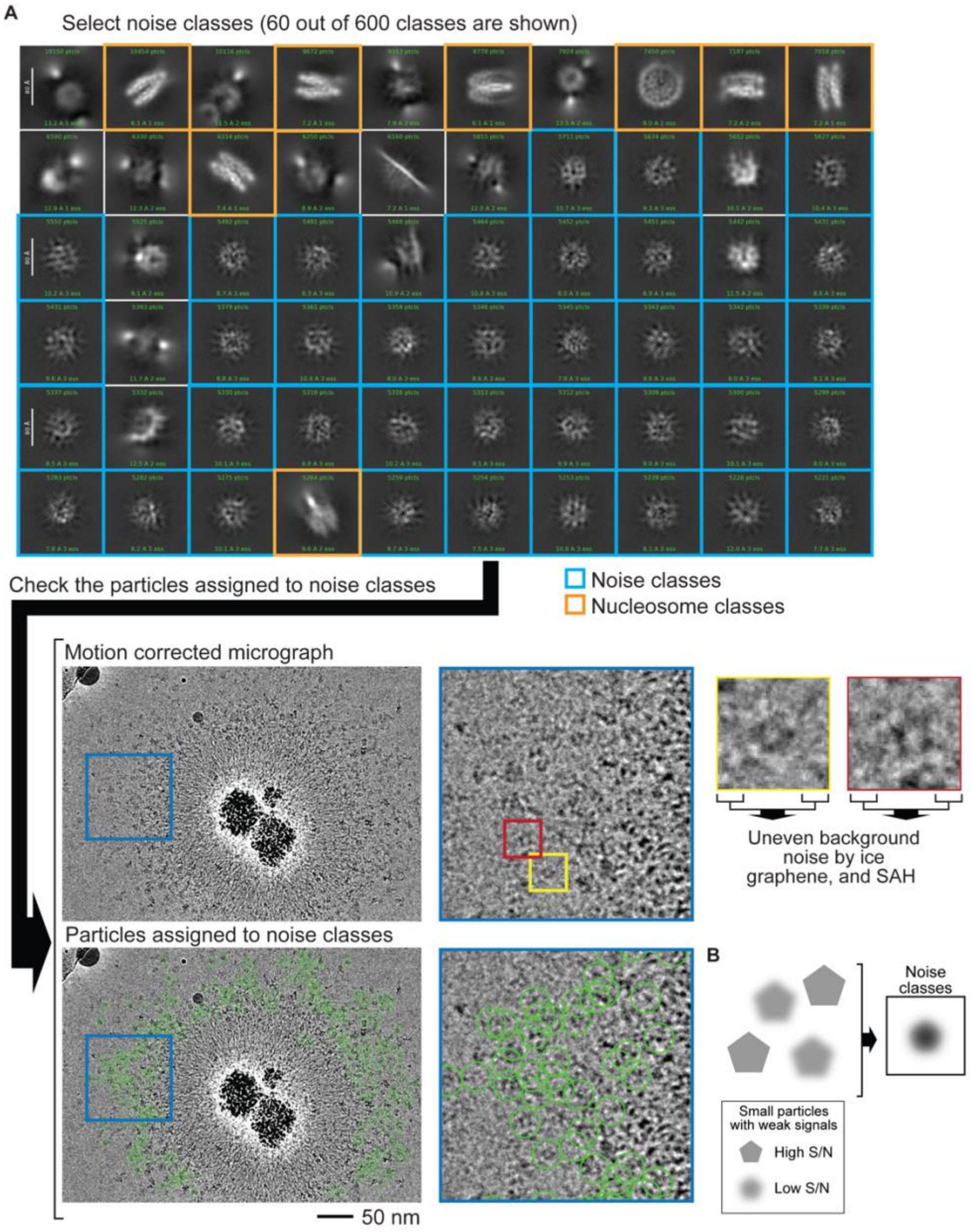

**Figure S6. The fraction with the interphase-specific GFP-H1.8 containing complex had many particles with low S/N by cryo-EM. (A)** The sucrose gradient fraction enriched with interphase-specific GFP-H1.8 containing complex (fraction 4 in Figure 3C) was subjected for MagIC-cryo-EM analysis. Initial 2D classification based on particles picked by Topaz generated only noise 2D classes (outlined with blue in the top panel) beside obvious nucleosome classes (outlined with orange in the top). Although 2D classes seem noisy, many of the original pick points marked apparent protein particles on the original motion-corrected micrograph (bottom). This suggests that these particle images were not properly aligned during 2D classification due to the low S/N of these particles. Uneven background noises were likely generated by uneven ice thickness, graphene and the SAH spacer proteins on MagIC-cryo-EM beads. **(B)** Graphical presentation of how noise 2D classes were generated. Although Topaz picked target protein particles on micrographs, many small target particles do not have strong enough S/N to be properly aligned during 2D classification.

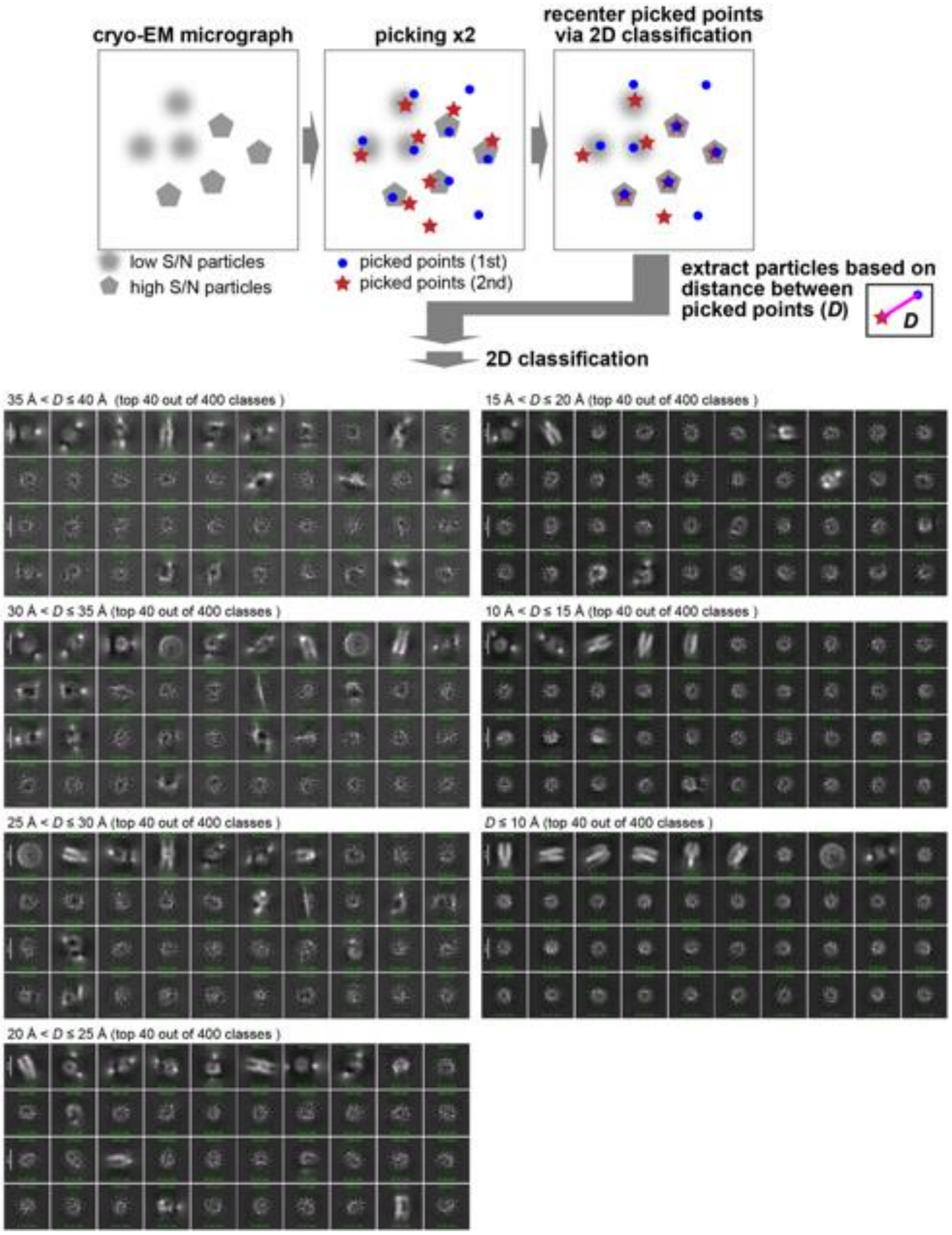

**Figure S7. Reproducible particle centering after 2D classification as a criterion for particles with high S/N.** The sucrose gradient fraction enriched with interphase-specific GFP-H1.8 containing complex (fraction 4 in Figure 3C) was isolated and analyzed by MagIC-cryo-EM. The initial particle locations assigned by particle picking software are updated during 2D to align the multiple images on a 2D map and place the reconstituted 2D map at the center of the reconstituted 2D space. To assess the reproducibility of particle centering during 2D classification, particle picking was repeated and subjected to the 2D classification individually. Particle images were sorted based on the distance *D* between a recentered picked point from the first picking set and another re-centered picked point from the second picking set. The sorted particle images were again applied to 2D classification. Particle images with *D* > 20 Å, which were not reproducibly recentered, generated noise 2D classes. In contrast, particle images with *D* ≤ 20 Å, which were reproducibly recentered, generated 2D classes with less background noises.

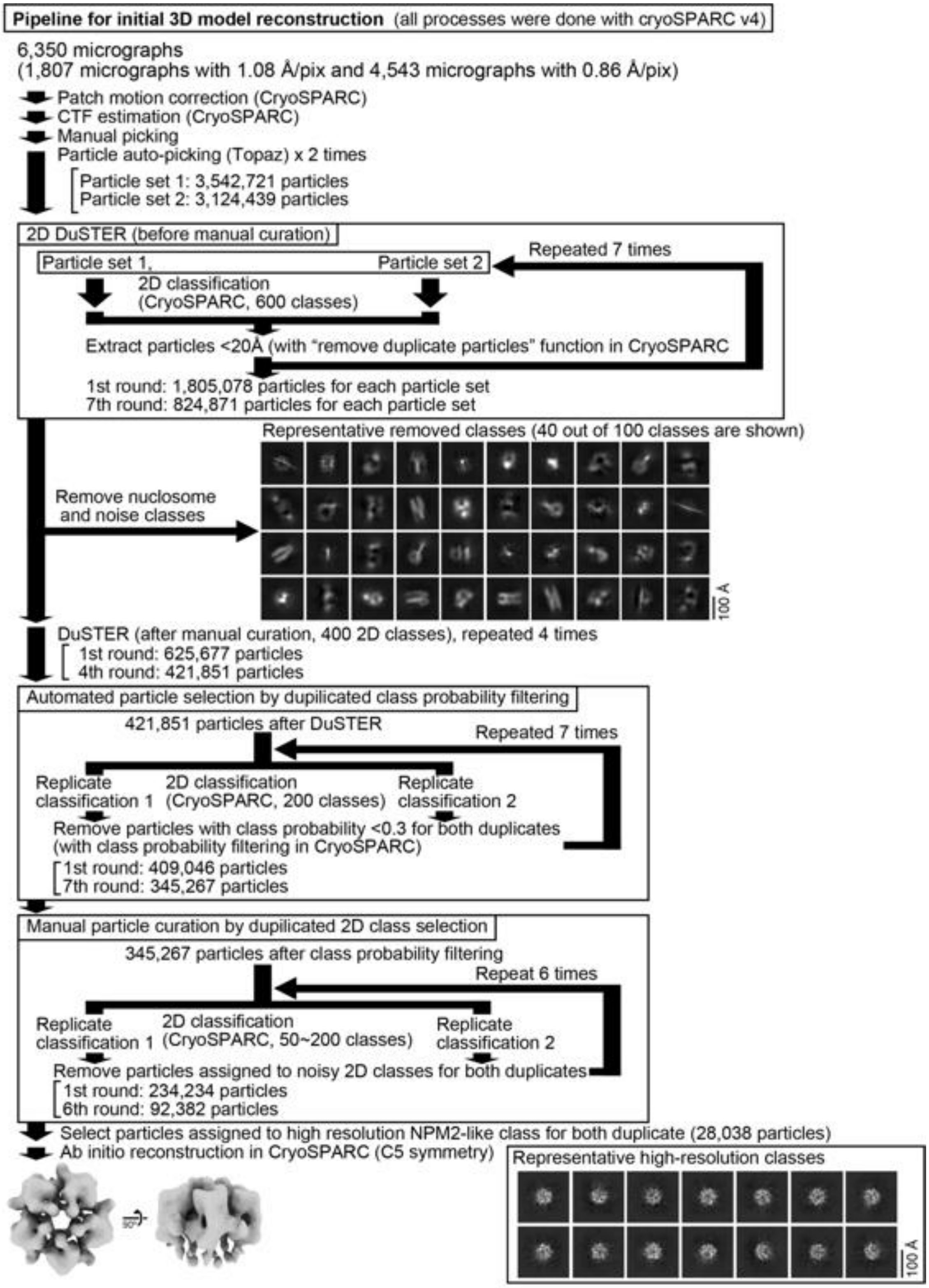

**Figure S8 Pipeline for 2D DuSTER for reconstructing a 3D initial model of the interphase-specific H1.8-containing complex (NPM2-H1.8) that is used as a template of 3D DuSTER.**

Please refer to the materials and methods section for a detailed description.

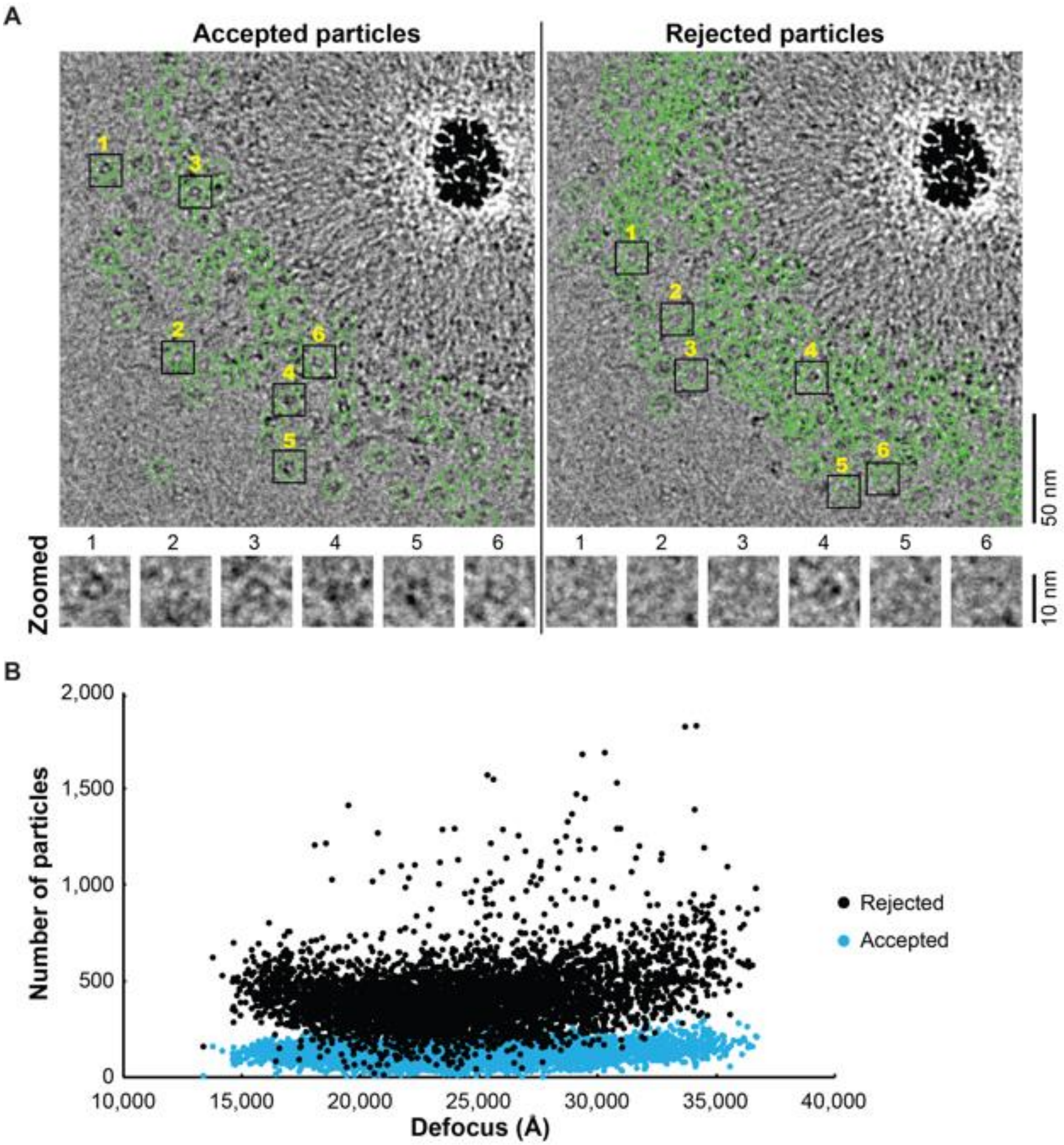

**Figure S9. The particles selected by DuSTER curation.**

**(A)** The particles accepted and rejected by DuSTER. Particles are mapped on motion-corrected micrographs. **(B)** DuSTER accepted particles across all defocus ranges. Numbers of accepted (cyan) and rejected (black) particles on each micrograph and their defocus length are plotted.

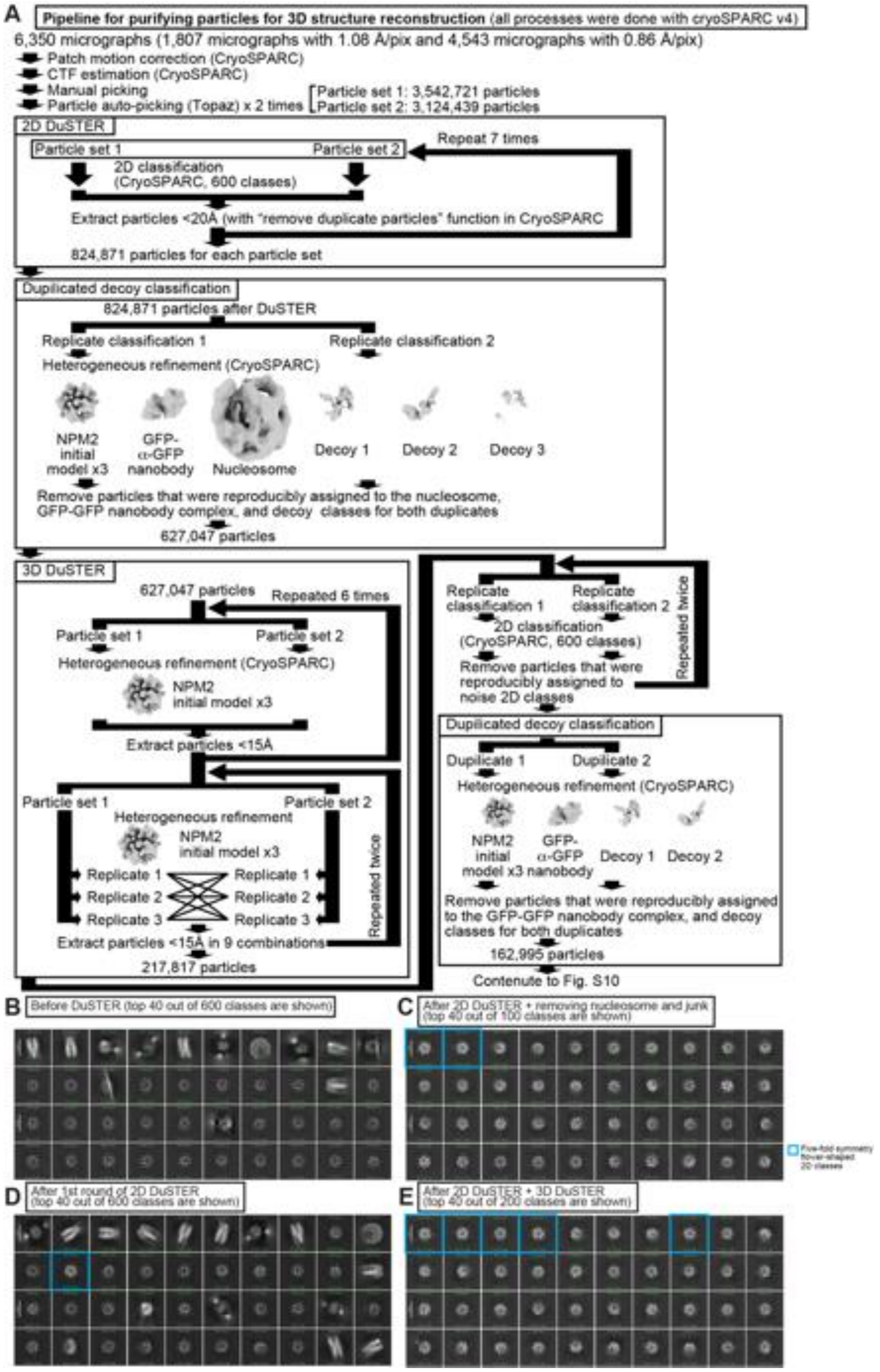

**Figure S10. Pipeline for 3D DuSTER.**

**(A)** Pipeline for 3D DuSTER. Please refer to the materials and methods section for a detailed description**. (B)** A 2D classification result of the particles picked by Topaz without particle curation with DuSTER. Beside obvious nucleosome classes, no reasonable 2D classes were observed before the DuSTER curation. **(C)** 2D classification of the particle after the single round of 2D DuSTER. Five-fold symmetry flower-shaped 2D classes (outlined with cyan) are observed. **(D)** 2D classification of the particles after seven rounds of 2D DuSTER and manual curation of non-target complex classes. **(E)** 2D classification of the particles after the completion of 2D DuSTER and 3D DuSTER.

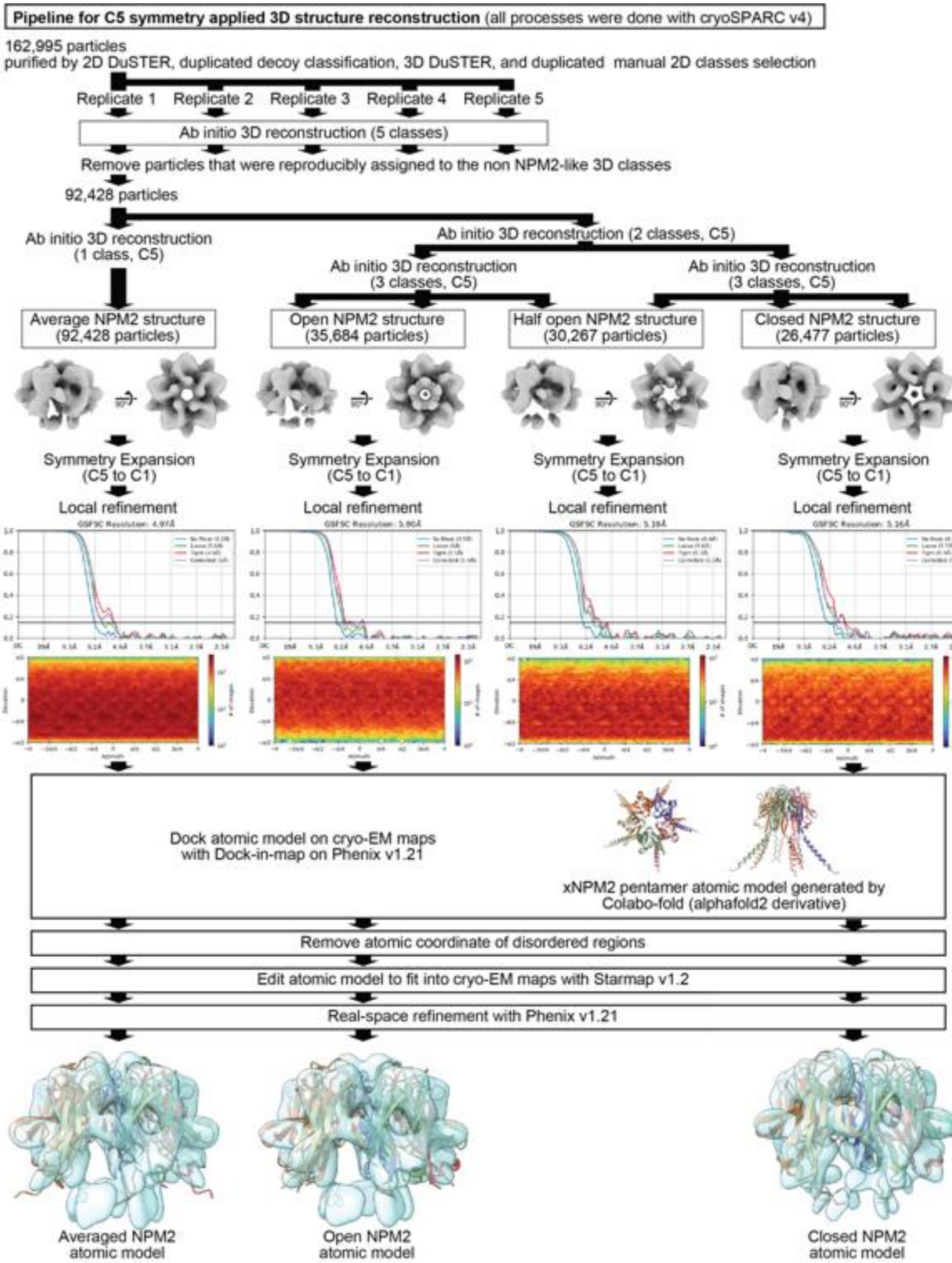

**Figure S11. Pipeline for 3D structure determination of the interphase-specific H1.8-containing complex (NPM2-H1.8).** Please refer to the materials and methods section for a detailed description**.**

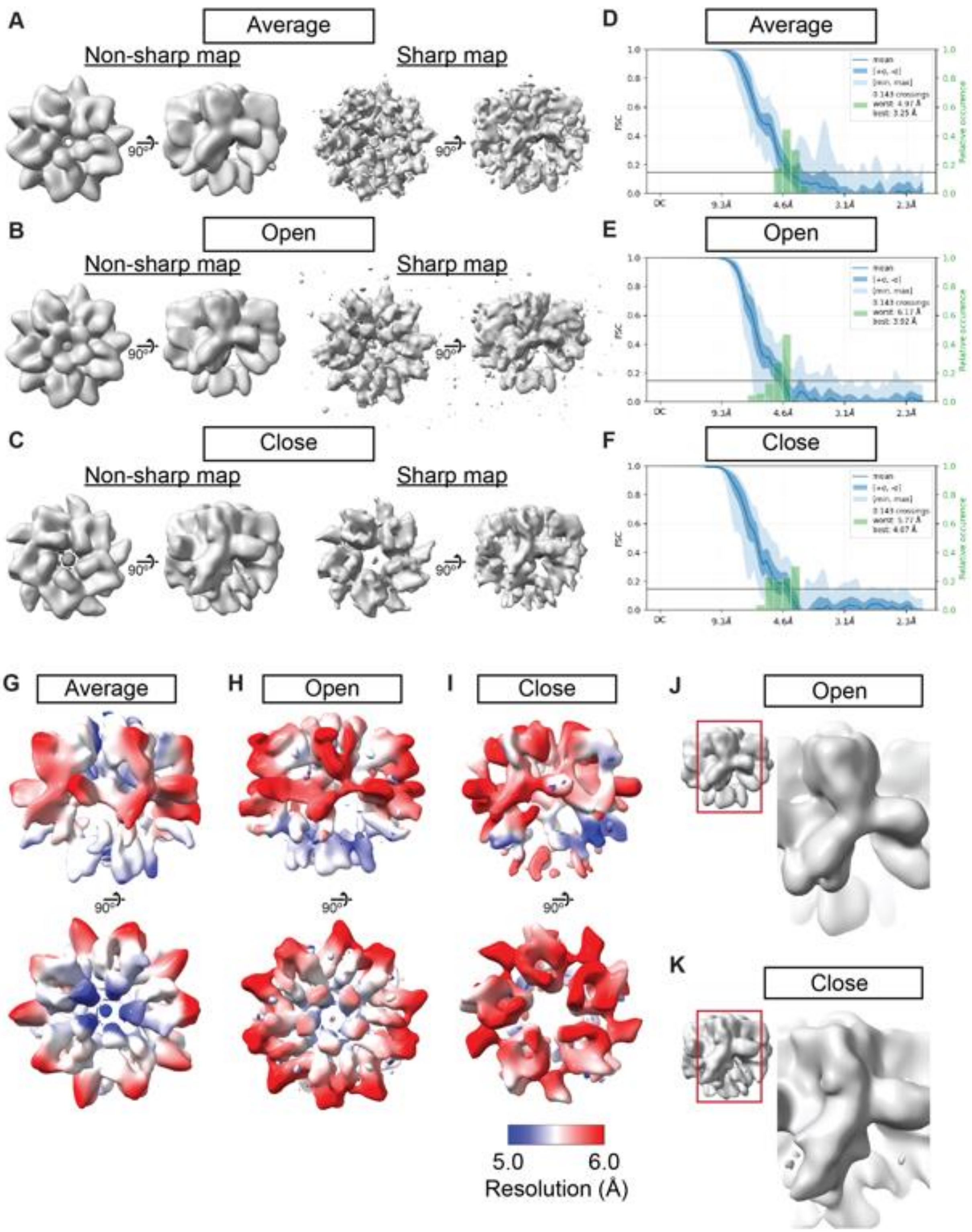

**Figure S12. Cryo-EM maps of NPM2 co-isolated with H1.8. (A-C)** The NPM2 cryo-EM maps before and after the map sharpening. **(D-F)** 3D FSC of the NPM2 cryo-EM maps **(G-I)** Local resolution of NPM2 cryo-EM maps. **(J-K)** The structural comparison of the open and closed form NPM2 cryo-EM maps.

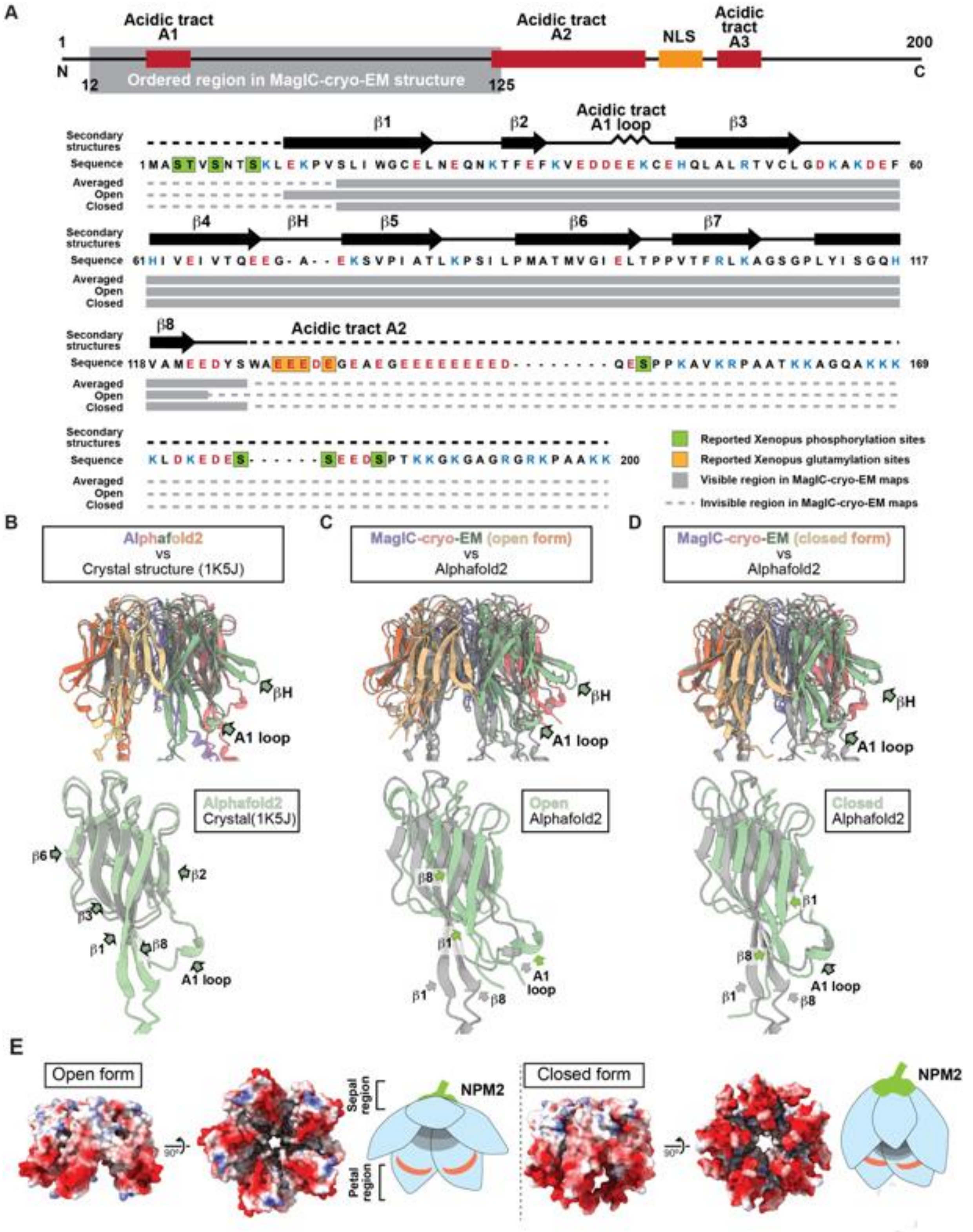

**Figure S13. Cryo-EM maps and atomic models of NPM2 co-isolated with H1.8. (A)** The structural comparison of the crystal structure of the pentameric NPM2 core (PDB ID: 1K5J), and AF2 predicted structure of the pentameric NPM2 core, and MagIC-cryo-EM structures of NPM2-H1.8. The MagIC-cryo-EM structures indicate NPM2 in the NPM2-H1.8 complex forms pentamer.

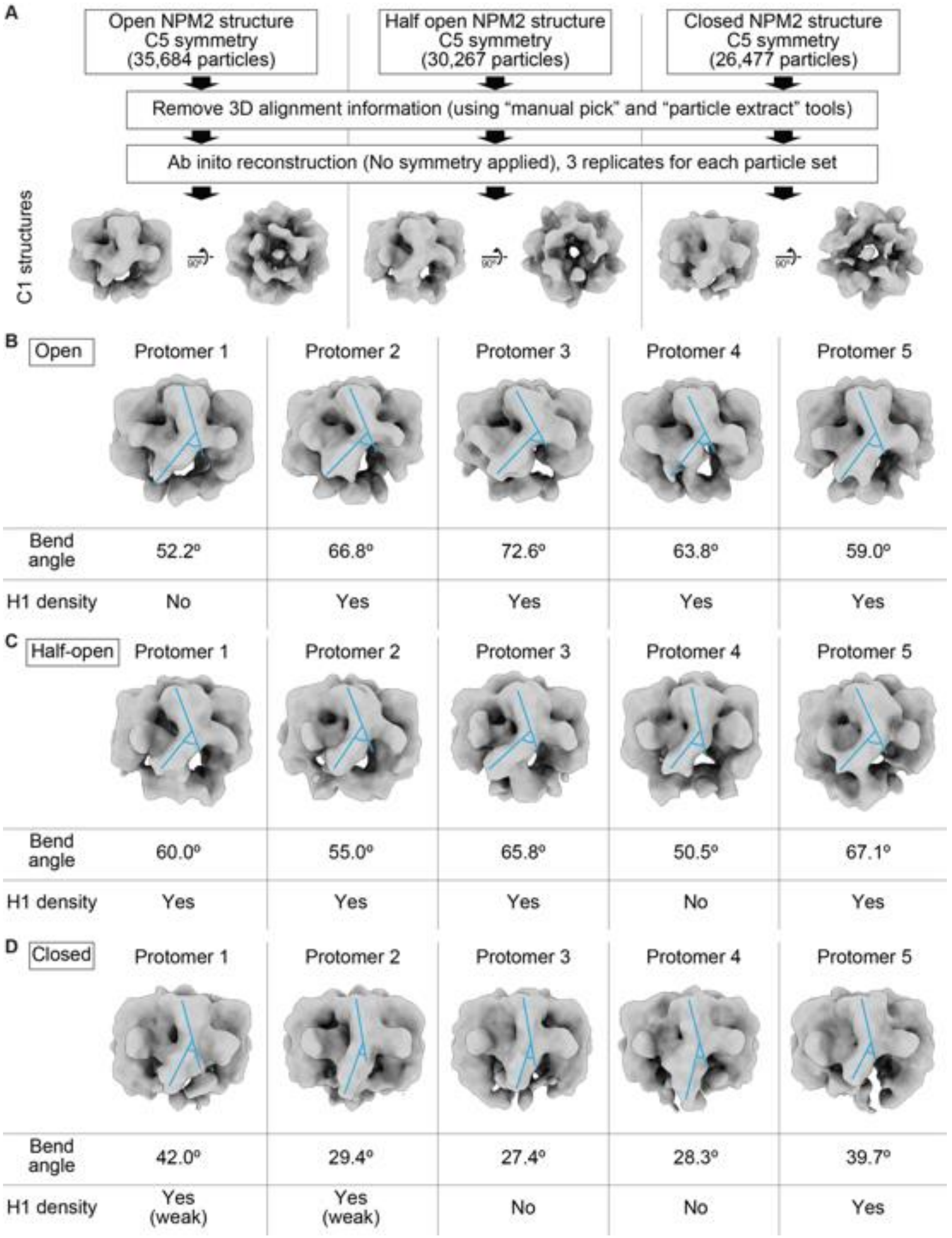

**Figure S14. Asymmetric structures of NPM2 co-isolated with H1.8 without applying C5 symmetry. (A)** Pipeline to reconstitute C1 symmetry NPM2-H1.8 structures. **(B-D)** Structural features of the open (B), half open (C), and closed (D) NPM2-H1.8 structures without applying C5 symmetry. Blue lines indicate the angles of the sepal and petal domains used for measuring the bend angles of each protomer. The bending is critical in forming the open-form NPM2 structures and increases the accessibility of the H1 binding sites. The bend angles and the bindings of the H1 densities are not consistent in each protomer, suggesting that NPM2 protomers with various openness co-exist in a single NPM2-H1.8 complex.

| Methods | Purity | Concentration | Volume | Amount | Advantage | Disadvantage |
|---|---|---|---|---|---|---|
| Cryo-EM (conventional) | Purified | 0.5 ~ 5.0 mg/mL | 3 ~ 4 μL/grid | > 10 μg/sample | simple | High concentration of sample is required |
| Cryo-EM (Jet vitrification) | Purified | 4 mg/mL | 0.001 μL/grid | >4 ng/sample | Very low volume of sample is required | High concentration of sample is required |
| Cryo-EM (Affinity grid) | Crude | 0.05 mg/mL | 3 ~ 4 μL/grid | > 1 μg/sample | Sample can be isolated and concentrated on grid | The maximum sample volume is limited |
| ChIP-seq | Crude | - | - | 10 ~ 50 ng DNA | Sample can be isolated and concentrated by beads | |
| SDS-PAGE (CBB stain) | Crude | 0.005 ~ 0.100 mg/mL | 1~20 μL/lane | > 30 ng/band | | |
| MagIC-cryo-EM | Crude | < 0.0005 mg/mL | 1 ~ 2000 μL | > 5 ng (2 ng DNA) /grid | Sample can be isolated and concentrated by beads | cryo-EM data collection points are selected manually. |
| SDS-PAGE (Silver stain) | Crude | 0.0001 ~ 0.001 mg/mL | 1~20 μL/lane | > 1 ng/band | | |

**Table S1. Sample requirements for the MagIC-cryo-EM and other biological approach.** To enable the structural analysis of the native protein complexes, we aimed to reduce the sample requirement of cryo-EM to that of ChIP-seq.

| Description | Full name or Functions | Input Interphase, fraction 5 | Input Metaphase, fraction 5 | MagIC-cryo-EM, Interphase, fraction 5 | MagIC-cryo-EM, Metaphase, fraction 5 |
|---|---|---|---|---|---|
| XBmRNA11963\| h2ac17.L | histone H2A | 1.37 x 10^10 | 4.12 x 10^9 | 2.08 x 10^8 | 1.49 x 10^8 |
| XBmRNA31731\| LOC121402261 | histone H2B | 7.87 x 10^9 | 2.26 x 10^9 | 1.10 x 10^8 | 2.25 x 10^8 |
| XBmRNA31368\| LOC121402047 | histone H4 | 7.34 x 10^9 | 2.00 x 10^9 | 1.11 x 10^8 | 1.63 x 10^8 |
| XBmRNA36987\| h2aj.L | histone H2A.J | 6.74 x 10^9 | 3.64 x 10^9 | 2.44 x 10^8 | 1.29 x 10^8 |
| XBmRNA58735\| h2ax.2.L | histone H2A.X | 6.56 x 10^9 | 1.73 x 10^9 | 2.44 x 10^8 | 1.75 x 10^8 |
| XBmRNA1949\| h2az1.L | histone H2A.Z | 4.50 x 10^9 | 3.64 x 10^9 | 2.44 x 10^8 | 1.52 x 10^8 |
| XBmRNA75391\| h3-3b.L | histone H3.3 | 3.33 x 10^9 | 1.19 x 10^9 | 3.84 x 10^7 | 5.29 x 10^7 |
| XBmRNA25971\| npm2.L | Nucleoplasmin | 1.77 x 10^9 | n.d. | 1.89 x 10^8 | n.d. |
| XBmRNA28658\| npm2.S | Nucleoplasmin | 1.76 x 10^9 | n.d. | 1.31 x 10^8 | n.d. |
| XBmRNA25690\| pcna.L | PCNA | 1.29 x 10^9 | n.d. | n.d. | n.d. |
| XBmRNA28883\| pcna.S | PCNA | 1.29 x 10^9 | n.d. | n.d. | n.d. |
| XBmRNA41314\| h1-8.S | Linker histone H1.8 | 7.12 x 10^8 | 1.93 x 10^8 | 5.67 x 10^7 | 6.87 x 10^7 |
| XBmRNA9392\| supt16h.S | supt16h (histone chaperone FACT complex component) | 3.14 x 10^8 | n.d. | n.d. | n.d. |
| XBmRNA4198\| supt16h.L | supt16h (histone chaperone FACT complex component) | 3.04 x 10^8 | n.d. | n.d. | n.d. |
| XBmRNA62381\| ssrp1.S | ssrp1 (histone chaperone FACT complex component) | 2.96 x 10^8 | n.d. | n.d. | n.d. |
| sfGFP | sfGFP (Tagged with H1.8) | 2.84 x 10^8 | 8.15 x 10^7 | 6.02 x 10^7 | 4.58 x 10^7 |
| XBmRNA26658\| dnmt1.L | DNA methyltransferase 1 | 1.41 x 10^8 | n.d. | n.d. | n.d. |
| XBmRNA31618\| dnmt1.S | DNA methyltransferase 1 | 1.35 x 10^8 | n.d. | n.d. | n.d. |
| XBmRNA5952\| sub1.L | Activated RNA polymerase II transcriptional coactivator p15 | 8.60 x 10^7 | n.d. | n.d. | n.d. |
| XBmRNA24726\| pclaf.L | PCNA-associated factor | 8.38 x 10^7 | n.d. | n.d. | n.d. |
| XBmRNA4881\| ran.L | GTP-binding nuclear protein Ran | 7.38 x 10^7 | n.d. | n.d. | n.d. |
| XBmRNA42021\| msh2.L | DNA repair protein MutS | 7.08 x 10^7 | n.d. | n.d. | n.d. |
| XBmRNA52420\| mcm4.L | DNA replication licensing factor MCM4 | 6.61 x 10^7 | n.d. | n.d. | n.d. |
| XBmRNA51197\| mcm6.L | DNA replication licensing factor MCM6 | 6.54 x 10^7 | n.d. | n.d. | n.d. |
| XBmRNA60815\| rcc2.L | Regulator of chromosome condensation 2 | 6.29 x 10^7 | n.d. | n.d. | n.d. |
| XBmRNA29645\| pclaf.S | PCNA-associated factor | 5.26 x 10^7 | n.d. | n.d. | n.d. |
| XBmRNA12919\| rcc1.L | Regulator of chromosome condensation 1 | 5.24 x 10^7 | n.d. | n.d. | n.d. |
| XBmRNA27060\| mcm7.L | DNA replication licensing factor MCM7 | 5.08 x 10^7 | n.d. | n.d. | n.d. |
| XBmRNA73621\| LOC108700788 | Importin subunit beta-like isoform X2 | 5.03 x 10^7 | n.d. | n.d. | n.d. |
| XBmRNA42017\| msh6.L | DNA repair protein MutS | 4.96 x 10^7 | n.d. | n.d. | n.d. |
| XBmRNA39469\| top1.2.S | DNA topoisomerase I | 4.92 x 10^7 | n.d. | n.d. | n.d. |
| XBmRNA82329\| kpna7.S | Importin subunit beta | 4.80 x 10^7 | n.d. | n.d. | n.d. |
| XBmRNA74418\| csnk2a1.L | Casein kinase II subunit alpha | 4.65 x 10^7 | n.d. | n.d. | n.d. |
| XBmRNA25938\| XB5867546.L | Nucleoplasmin isoform (lacking C-ter tail) | 4.31 x 10^7 | n.d. | n.d. | n.d. |
| XBmRNA7871\| uhrf1.S | E3 ubiquitin-protein ligase UHRF1 | 4.27 x 10^7 | n.d. | n.d. | n.d. |
| XBmRNA12245\| rpa1.L | Replication protein A | 4.10 x 10^7 | n.d. | n.d. | n.d. |
| XBmRNA41384\| LOC100192369 | Sperm-specific nuclear basic protein 1 | 4.08 x 10^7 | n.d. | n.d. | n.d. |
| XBmRNA52377\| mcm3.L | DNA replication licensing factor MCM3 | 4.04 x 10^7 | n.d. | n.d. | n.d. |
| XBmRNA36255\| mcm5.L | DNA replication licensing factor MCM5 | 3.86 x 10^7 | n.d. | n.d. | n.d. |
| XBmRNA36962\| mcm2.L | DNA replication licensing factor MCM2 | 3.73 x 10^7 | n.d. | n.d. | n.d. |
| XBmRNA60207\| pold1.L | DNA polymerase | 3.63 x 10^7 | n.d. | n.d. | n.d. |
| XBmRNA33152\| fen1.L | Flap endonuclease 1-A | 3.59 x 10^7 | n.d. | n.d. | n.d. |
| XBmRNA25831\| pold2.L | DNA polymerase delta subunit 2 | 3.37 x 10^7 | n.d. | n.d. | n.d. |
| XBmRNA38234\| ddb1.S | DNA damage-binding protein 1 | 2.98 x 10^7 | n.d. | n.d. | n.d. |
| XBmRNA78857\| dppa2.L | Developmental pluripotency associated 2 | 2.96 x 10^7 | n.d. | n.d. | n.d. |
| XBmRNA34892\| gins3.L | DNA replication complex GINS protein PSF3 | 2.93 x 10^7 | n.d. | n.d. | n.d. |
| XBmRNA83211\| hirip3.S | HIRA-interacting protein 3 | 2.69 x 10^7 | n.d. | n.d. | n.d. |
| XBmRNA35380\| nasp.L | histone chaperone NASP | 1.84 x 10^7 | n.d. | n.d. | n.d. |
| XBmRNA23497\| hmga2.L | High mobility group AT-hook 2 | 1.45 x 10^7 | n.d. | n.d. | n.d. |
| XBmRNA30566\| hmga2.S | High mobility group AT-hook 2 | 1.43 x 10^7 | n.d. | n.d. | n.d. |
| XBmRNA33324\| pola2.L | DNA polymerase alpha subunit B | 1.10 x 10^7 | n.d. | n.d. | n.d. |
| Streptavidin | MagIC-cryo-EM beads proteins | n.d. | n.d. | 1.22 x 10^9 | 2.19 x 10^9 |
| SPY-tagGFPnanobody | MagIC-cryo-EM beads proteins | n.d. | n.d. | 7.69 x 10^8 | 1.41 x 10^9 |
| Spycatcher3 | MagIC-cryo-EM beads proteins | n.d. | n.d. | 5.99 x 10^8 | 7.82 x 10^8 |
| 11nm_3HB | MagIC-cryo-EM beads proteins | n.d. | n.d. | 4.32 x 10^7 | 5.90 x 10^7 |
| 60nm_SAH | MagIC-cryo-EM beads proteins | n.d. | n.d. | 1.21 x 10^7 | 7.73 x 10^6 |

**Table S2. The list of the chromatin proteins detected by mass spectrometry before and after enrichment on the MagIC-cryo-EM-beads.** The MS analysis was conducted to the sucrose gradient fractions 5, before (Figure 3C) and after (Figure 3E) enrichment with the GPF nanobody-MagIC-cryo-EM beads. Detectable MS signals for known chromatin proteins and the recombinant proteins used for assembling the MagIC-cryo-EM beads were manually selected and are listed here. See table S5 of the full MS data.

Table S3.1 Molecular wight of the components

| Name | Molecular wight (Da) |
|---|---|
| NPM2 monomer | 21,917 |
| NPM2 pentamer | 109,587 |
| NPM2 decamer | 219,175 |
| H1.8-GFP | 56,704 |

Table S3.2 Molecular wight of the complexes

| Name | Molecular wight (Da) |
|---|---|
| NPM2 pentamer + H1.8-GFP monomer | 166,291 |
| NPM2 pentamer + H1.8-GFP pentamer | 393,107 |
| NPM2 decamer + H1.8-GFP monomer | 275,878 |
| NPM2 decamer + H1.8-GFP pentamer | 502,694 |
| Nucleosome (193bp DNA) | 228,111 |

**Table S3.** Expected mass of the NPM2-H1.8-GFP complex. Sucrose gradient elution volume indicates that the NPM2-H1.8-GFP complex is smaller than mono-nucleosome (around 230 kDa). Only the NMP2 pentamer complexed H1.8-GFP monomer (166 kDa) reasonably explains the sucrose gradient result.

| Sample name | Nucleosome in polynucleosome attached on magnetic beads | *in vitro* reconstituted H1.8-GFP nucleosome MagIC-cryo-EM | Interphase H1.8-GFP nucleosome MagIC-cryo-EM | Metaphase H1.8-GFP nucleosome MagIC-cryo-EM | Interphase & metaphase mixed H1.8-GFP nucleosome MagIC-cryo-EM | Interphase H1.8-GFP-NPM2 MagIC-cryo-EM | | |
|---|---|---|---|---|---|---|---|---|
| Deta collection | | | | | | | | |
| Microscope | Talos Arctica | Titan Krios | Titan Krios | Titan Krios | - | Titan Krios | | |
| Magnification | 25,000 | 53,000 | 53,000 | 53,000 | - | 64,000 / 81,000 | | |
| Voltage (kV) | 200 | 300 | 300 | 300 | - | 300 | | |
| Camera | Gatan K2 Summit | Gatan K3 | Gatan K3 | Gatan K3 | - | Gatan K3 | | |
| Electron exposure (e–/Å) | 34.4 | 45.45 | 45.45 | 45.45 | - | 45.3 / 51.92 | | |
| Defocus range (μm) | -1.5 ~ -2.5 | -2 ~ -3.5 | -2 ~ -3.5 | -2 ~ -3.5 | - | -2.0 ~ -3.5 / -1.5 ~ -3.5 | | |
| Pixel size (Å) | 1.5 | 1.32 | 1.32 | 1.32 | - | 1.08 / 0.86 | | |
| Micrographs | 734 | 1,890 | 677 | 965 | - | 1,807 / 4,543 | | |
| Data prosessing | | | | | | Avaraged | Open | Closed |
| Particle images (no.) | 41,000 | 184,706 | 20,639 | 22,051 | 42,690 | 92428 | 35684 | 26477 |
| Symmetry imposed | C1 | C1 | C1 | C1 | C1 | C5 | C5 | C5 |
| Map resolution (FSC=0.143) | 4.8 | 3.6 | 4.3 | 4.1 | 3.9 | 5.0 | 5.9 | 5.2 |
| EMDR ID | EMD-42599 | EMD-42598 | EMD-42596 | EMD-42597 | EMD-42594 | EMD-43238 | EMD-43239 | EMD-43240 |
| Atomic models | | | | | | Avaraged | Open | Closed |
| Chains | | | | | | 5 | 5 | 5 |
| Atoms | | | | | | 8535 (Hydrogens: 4245) | 4315 (Hydrogens: 0) | 4290 (Hydrogens: 0) |
| Residues | | | | | | Protein: 550 Nucleotide: 0 | Protein: 555 Nucleotide: 0 | Protein: 550 Nucleotide: 0 |
| Water | | | | | | 0 | 0 | 0 |
| Ligands | | | | | | 0 | 0 | 0 |
| Bonds (RMSD) | | | | | | | | |
| Length (Å) (# > 4σ) | | | | | | 0.006 (15) | 0.003 (0) | 0.003 (0) |
| Angles (°) (# > 4σ) | | | | | | 0.699 (10) | 0.758 (5) | 0.884 (0) |
| MolProbity score | | | | | | 1.98 | 2.22 | 2.16 |
| Clash score | | | | | | 14.32 | 22.71 | 26.17 |
| Ramachandran plot (%) | | | | | | | | |
| Outliers | | | | | | 0 | 0 | 0 |
| Allowed | | | | | | 4.63 | 5.5 | 3.7 |
| Favored | | | | | | 95.37 | 94.5 | 96.3 |
| Rama-Z (Ramachandran plot Z-score, RMSD) | | | | | | | | |
| whole (N = 540) | | | | | | -2.46 (0.30) | -2.90 (0.29) | -1.28 (0.33) |
| helix (N = 0) | | | | | | --- (---) | -3.48 (0.84) | -1.84 (1.07) |
| sheet (N = 160) | | | | | | -1.74 (0.46) | -1.27 (0.35) | -0.38 (0.31) |
| loop (N = 380) | | | | | | -1.71 (0.24) | -2.21 (0.26) | -1.10 (0.33) |
| Rotamer outliers (%) | | | | | | 0 | 0 | 0 |
| Cβ outliers (%) | | | | | | NA | NA | NA |
| Peptide plane (%) | | | | | | | | |
| Cis proline/general | | | | | | 33.3/0.0 | 28.6/0.0 | 33.3/0.0 |
| Twisted proline/general | | | | | | 0.0/0.0 | 0.0/0.0 | 0.0/1.0 |
| CaBLAM outliers (%) | | | | | | 2.83 | 1.87 | 3.77 |
| ADP (B-factors) | | | | | | | | |
| Iso/Aniso (#) | | | | | | 4290/0 | 4315/0 | 4290/0 |
| min/max/mean | | | | | | | | |
| Protein | | | | | | 311.84/446.13/375.67 | 250.02/795.91/470.81 | 343.54/628.84/480.26 |
| Nucleotide | | | | | | --- | --- | --- |
| Ligand | | | | | | --- | --- | --- |
| Water | | | | | | --- | --- | --- |
| Occupancy | | | | | | | | |
| Mean | | | | | | 1 | 1 | 1 |
| occ = 1 (%) | | | | | | 100 | 100 | 100 |
| 0 < occ < 1 (%) | | | | | | 0 | 0 | 0 |
| occ > 1 (%) | | | | | | 0 | 0 | 0 |
| Lengths (Å) | | | | | | 82.08, 83.16, 62.64 | 84.24, 85.32, 58.32 | 79.92, 81.00, 69.12 |
| Angles (°) | | | | | | 90.00, 90.00, 90.00 | 90.00, 90.00, 90.00 | 90.00, 90.00, 90.00 |
| Supplied Resolution (Å) | | | | | | 6 | 6 | 6 |
| Resolution Estimates (Å) | | | | | | Masked // Unmasked | Masked // Unmasked | Masked // Unmasked |
| d FSC (half maps; 0.143) | | | | | | --- // --- | --- // --- | --- // --- |
| d 99 (full/half1/half2) | | | | | | 8.5/---/--- // 8.5/---/--- | 8.1/---/--- // 8.1/---/--- | 8.3/---/--- // 8.3/---/--- |
| d model | | | | | | 7.7 // 7.7 | 8.5 // 8.5 | 7.7 // 7.7 |
| d FSC model (0/0.143/0.5) | | | | | | 4.3/6.2/18.9 // 4.3/6.2/2.6 | 5.5/6.3/18.2 // 5.5/6.3/18.2 | 5.0/6.1/12.2 // 5.0/6.1/11.3 |
| Map min/max/mean | | | | | | -9.459459459 | -10.46511628 | -15.15151515 |
| CC (mask) | | | | | | 0.67 | 0.67 | 0.69 |
| CC (box) | | | | | | 0.79 | 0.75 | 0.8 |
| CC (peaks) | | | | | | 0.56 | 0.56 | 0.62 |
| CC (volume) | | | | | | 0.66 | 0.67 | 0.69 |
| Mean CC for ligands | | | | | | --- | --- | --- |
| PDB ID | | | | | | 8VHI | 8VHJ | 8VHK |

**Table S4** Statistics of the cryo-EM structures.
